# Paracrine control of α-cell glucagon exocytosis is compromised in human type-2 diabetes

**DOI:** 10.1101/789545

**Authors:** Muhmmad Omar-Hmeadi, Per-Eric Lund, Nikhil R Gandasi, Anders Tengholm, Sebastian Barg

## Abstract

Glucagon is secreted from pancreatic α-cells to activate gluconeogenesis and other pathways that raise blood glucose during hypoglycemia. Glucose-dependent regulation of glucagon secretion involves both α-cell-intrinsic mechanisms and paracrine control through insulin and somatostatin. In type-2 diabetes (T2D) inadequately high glucagon levels contribute to hyperglycemia. To understand these disease-associated changes at the cellular level, and to isolate intrinsic and paracrine effects, we analyzed glucagon granule exocytosis and membrane excitability in isolated α-cells from 56 non-diabetic (ND) and 15 T2D human donors. High resolution imaging showed that glucagon granule exocytosis had a U-shaped sensitivity to glucose, with the slowest rate around 7 mM glucose, and accelerated rates at <5 and >10 mM glucose. Exocytosis was reduced in T2D α-cells, but their glucose sensitivity remained intact and there were no changes in voltage-dependent ion currents or granule trafficking. Instead, α-cells from T2D donors were markedly insensitive to somatostatin and insulin, which rapidly inhibited exocytosis and electrical activity in ND cells. Thus, intrinsic mechanisms do not inhibit glucagon secretion at hyperglycemia, and elevated glucagon levels in human T2D reflect an insensitivity of α-cells to paracrine inhibition.

## Introduction

Glucagon is released from pancreatic α-cells and counteracts the glucose-lowering actions of insulin by stimulating gluconeogenesis and hepatic glucose output. Initially thought of only as part of the body’s defense against hypoglycemia ^1^, it is now clear that inadequate glucagon levels also contribute to diabetic hyperglycemia and present a challenge for diabetes management ^2,3^. Glucagon secretion is triggered by low blood glucose and suppressed at physiological glucose levels, and both α-cell intrinsic and paracrine mechanisms have been cited to explain these effects. In the intrinsic models, glucose metabolism and the generation of ATP play a central role ^4–6^, either through subtle, K_ATP_ channel-dependent depolarization of the resting membrane potential and subsequent inactivation of Na^+^-channels ^7–10^, or as consequence of glucose-induced activation of the sarco/endoplasmic reticulum Ca^2+^-ATPase that leads to closure of store-operated channels and hyperpolarization ^11,12^. In addition, intrinsic glucose-dependent cAMP signaling may play a role ^13^. However, none of these models fully explain the glucose concentration dependence of glucagon secretion, in particular in the hyperglycemic range.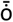

Glucagon secretion is also under paracrine control from neighboring β- and δ-cells, and the inhibitory effects of somatostatin ^14–18^, insulin ^19–23^, and GABA ^24,25^ have long been recognized. Paracrine inhibition is likely to play a role at elevated glucose levels, when β- and δ-cells are active. Indeed, glucagon is secreted in pulses that are anti-synchronous to pulses of insulin and somatostatin ^26,27^. This relationship is important for the postprandial suppression of glucagon secretion and is lost in type-2 diabetes and pre-diabetes ^28–30^. Human and rat α-cells express the somatostatin receptor SSTR2, which leads to hyperpolarization via activation of GIRK-channels ^31,32^. At least in rats, somatostatin also inhibits the exocytosis machinery via calcineurin ^32^, and inhibits α-cell exocytosis by effectively decreasing cytosolic cAMP ^33,34^. Insulin receptor signaling is required for the suppression of glucagon secretion in vivo ^35^, but the precise mechanisms behind this are still debated ^34,36,37^. Decreased sensitivity to insulin (or somatostatin) may therefore underlie the inadequate glucagon secretion in type-2 diabetes ^38^.

Glucagon is stored in ∼7000 granules (diameter ∼270 nm) and secreted by Ca^2+^- and SNARE protein-dependent exocytosis ^39,40^. At any time, only about 1% of these granules are in a releasable state that can undergo exocytosis upon Ca^2+^-influx ^41^. Paracrine signaling and glucose regulate glucagon secretion at least in part by affecting the size of this releasable pool of granules ^32,42,43^. In many endocrine cells, secretory granules become release ready by sequential docking at the plasma membrane and assembly of the secretory machinery (priming) ^44,45^. Although disturbances in these steps have been documented in β-cells of type-2 diabetic donors ^46–49^, they have not been studied in α-cells.

An obstacle for understanding the regulation of α-cells has been the difficulty to isolate intrinsic and paracrine factors of α-cell regulation, as well as species differences between humans and rodent models. Glucagon secretion in vivo and in intact islets is affected by the presence of neighboring cell types, while single cell electrophysiological measurements are invasive and may not reflect the in vivo situation. In the current work, we took an optical approach to study glucagon granule exocytosis in isolated α-cells of 56 non-diabetic and 15 type-2 diabetic human subjects. We report a U-shaped exocytosis response to glucose that was somewhat reduced in T2D α-cells, due to decreased availability of releasable granules, rather than granule docking. Importantly, paracrine inhibition of exocytosis and electrical activity by insulin and somatostatin was strongly impaired in T2D α-cells, which might underlie the hyperglucagonemia in type-2 diabetes.

## Results

### Exocytosis of glucagon granules in pancreatic α-cells

Docking and exocytosis of glucagon granules at the plasma membrane was studied in dispersed islet preparations from 56 non-diabetic (ND) donors with glycated hemoglobin (HbA1c) values <6% (average 5.57±0.29%, table S1). To identify α-cells, we transduced with Pppg-EGFP and the secretory granule marker NPY-mCherry, or with Pppg-NPY-EGFP (Fig 1A and S1A,B) to drive expression of fluorescent proteins from the pre-proglucagon promoter (see methods). After culture for 26-48 h, the cells were imaged by total internal reflection (TIRF) microscopy, which selectively images fluorescence near the plasma membrane (exponential decay constant τ ∼0.1 µm). The granule marker displayed a punctate staining pattern and excellent overlap with anti-glucagon immunostaining (Fig 1A). Local application of elevated K^+^ (75 mM, replacing Na^+^) to depolarize the cells resulted in exocytosis, seen as rapid disappearance of some of the individual fluorescently labeled granules (Fig 1B and examples in S1B,C). Most granules within the field and all granules undergoing exocytosis were essentially immobile during the experiment, suggesting that they were molecularly tethered or docked at the plasma membrane. We pooled data from ND α-cells (n=158 cells from 26 donors), which showed that 0.077±0.004 gr*;µm^-2^ or 12±0.8% of the docked granules were released during short (40s) exposure to elevated K^+^ (Fig 1C, black). Exocytosis proceeded initially at high rate (5.2*;10^−3^ gr*;s^-1^*;µm^-2^ during the first 10 s of stimulation) and decreased later to less than 0.6*;10^−3^ gr*;s^-1^*;µm^-2^; these rates are only about one-third of those observed in human β-cells ^46^. Fitting with a double exponential function revealed a fast (τ=3.6±0.2 s; 39±3%; n=1325 granules) and a slow component (19.9±0.9 s; Fig 1C). We also noticed that stimulation with K^+^ resulted in partial depletion of docked granules (Fig 1D), suggesting that docking of granules to replace those lost by exocytosis is a relatively slow process. This notion was confirmed by using a double stimulation protocol (Fig S2). Thus, exocytosis of glucagon granules occurs with a biphasic pattern that is reminiscent of the exocytosis kinetics in other endocrine cells, suggesting the existence of granule pools with differing release probability.

**Fig 1.**
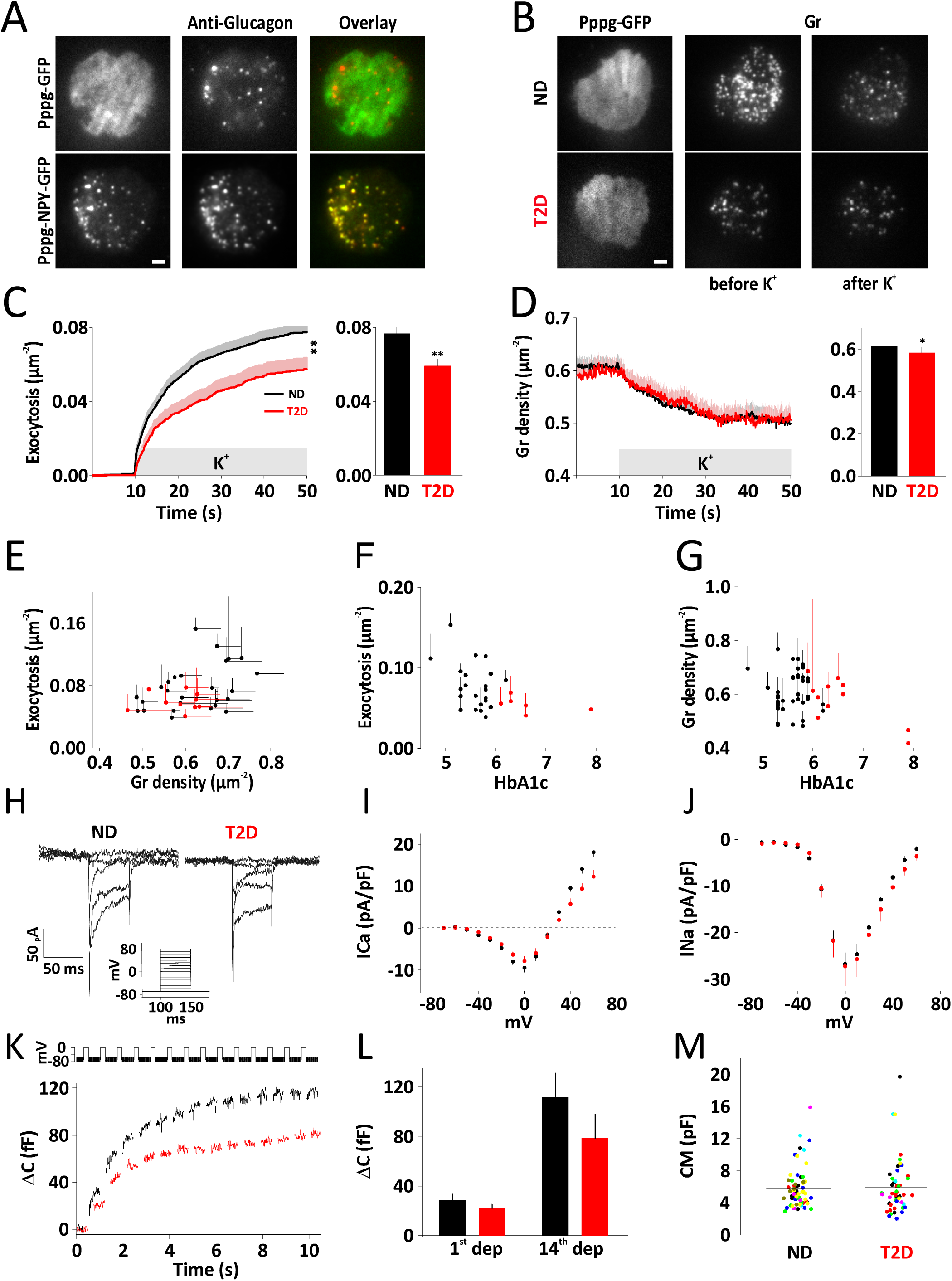
Exocytosis of glucagon granules in normal and diabetic pancreatic α-cells. **(A)** TIRF images of dispersed human islet cells transduced with Pppg-EGFP (top) or Pppg-NPY-EGFP (bottom). 90 % of EGFP expressing cells (n= 91 cells, 5 donors) and 93% of NPY-EGFP expressing cells (n=70 cells, 4 donors) were positive for glucagon. Scale bar 2 µm. **(B)** Examples of cells from non-diabetic (ND) and type 2 diabetic (T2D) donors expressing Pppg-EGFP together with granule marker (gr) or expressing Pppg-NPY-EGFP(not shown). The cells were imaged by TIRF microscopy and examples are before and after stimulation with 75mM K^+^ for 40s. (K^+^ was elevated during 10-50 s). Scale bar 2 µm. **(C)** Average cumulative number of exocytotic events in experiments as in (B) plotted over time and normalized to the cell area (left, cell wise); 1325 granules in 152 ND cells (black), 389 granules in 63 T2D cells (red). Exocytosis (right, donor wise) was 0.059±0.003 gr*;µm-2 in 10 T2D donors compared with 0.076±0.005 gr*;µm-2 in 26 ND donors. *;*; indicates p<0.01 (t-test). **(D)** Time courses of granule (gr) density (left, cell wise) in ND or T2D cells as in (B). Glucagon density (right, donor wise) was 0.58±0.027 gr*;µm-2 in 15 T2D donors compared with 0.61±0.003 gr*;µm-2 in 45 ND donors. *; indicates p<0.05 (t-test). **(E)** Exocytosis after stimulation with 75mM K^+^ for 40s as function of granule density during K^+^ stimulation. Each symbol in (E, F, G) represents represent individual donors ± SEM (5-20 cells for each donor; ND donors are shown in black, T2D in red). Correlation was quantified as Pearson coefficient r using Excel. **(F)** Exocytosis after stimulation with 75mM K^+^ for 40s as function of donor HbA1c. **(G)** Granule density as function of donor HbA1c. **(H)** Ca^2+^ and Na^+^-currents raw current figure from human non-diabetic and diabetic α-cells. Currents were elicited by a 50 ms depolarizing pulses (−70 to +80 mV in 10 mV increments) from a holding potential of −70 mV. For clarity, only the responses between –40 mV and +10 are shown. **(I-J)** Current (I)-voltage (V) relationships for Ca2+ and Na+ currents recorded from ND (n=38, 4 donors, black) and diabetic T2D (n=32, 3 donors, red) human pancreatic a-cells. Currents have been normalized to cell size (pF). **(K-L)** Cell capacitance during a train consisting of 14 200-ms depolarization from −70 mV to 0 mV (stimulation frequency: 1.3 Hz). **(L)** Average Change in membrane capacitance (ΔCm) in response to the 1st and the total increase during the entire train of 14 pulses (Σ1-14), from ND (n=20, 4 donors, black) and diabetic T2D (n=18, 3 donors, red) human pancreatic α-cells. **(M)** Differences in cell size between T2D (n=48, 7 donors, red) and ND (n=66, black, 8 donors, black) human α-cells. Line represents mean values.

### Exocytosis of glucagon granules in diabetic pancreatic α-cells

During the course of this study, we also received islets from 15 donors that had been diagnosed clinically with type-2 diabetes (T2D), or whose glycated hemoglobin (HbA1c) values were above 6% (average 6.6±0.7%, table S1). When α-cells from these T2D donors were stimulated by depolarization with high K^+^ (Fig 1B), the rate of exocytosis was less than that observed in ND α-cells (0.057±0.006 gr*;µm^-2^ or 75±10% of that observed in ND α-cells; n=63 T2D cells from 10 donors, p=0.004; Fig 1C). This was mostly due to a reduced proportion of the fast component (τ=2.1±0.1 s, 25±3%, n=389 granules, p=0.0008 vs ND). Since all exocytosis originated from granules that had been resident at the plasma membrane prior to stimulation, we also quantified docked granules. The density of granules in TIRF images was slightly reduced in T2D α-cells (0.57±0.02 gr*;µm^-2^; n=94 T2D cells from 15 donors) compared with ND (0.61±0.008 gr*;µm^-2^; n=382 ND cells from 45 donors, Fig 1D). The finding that donor averages of K^+^-stimulated exocytosis and granule density correlated (Pearson r=0.39, p=0.01, 36 donors; Fig 1E) suggests that impaired granule docking underlies the reduced capacity for exocytosis, which is similar to what we have previously reported for human β-cells ^46^. Interestingly, both K^+^-stimulated exocytosis (r=0.46, p=0.01; Fig 1F) and docked granules (r=0.31, p=0.02; Fig 1G) anti-correlated with the donor’s HbA1c. The relationships are surprising given that reduced glucagon levels should lead to reduced blood glucose and HbA1c values, and suggest that the diabetic state is causal for the reduced release capacity of α-cells.

Since exocytosis in α-cells depends on Ca^2+^-influx, we used patch clamp electrophysiology to measure voltage-dependent ion currents. ND or T2D cells were voltage clamped in whole-cell mode, and subjected to step depolarizations up to +70 mV from a holding potential of −70 mV (Fig 1H). Analysis of the resulting inward currents revealed peak Ca^2+^ (Fig 1I) and Na^+^-currents (Fig 1J) that were of similar amplitude in ND and T2D cells. Half-maximal Ca^2+^-current activation was reached at −23±0.4 (ND) and −25±1.4 mV (T2D, n.s.), and half-maximal Na^+^-current activation was at −25±0.6 (ND) and −24±0.8 mV (T2D, n.s.). We also determined depolarization evoked membrane capacitance increases as measure of exocytosis. A train of 14 depolarizations to zero mV lasting 200 ms each resulted in a total capacitance increase of 112±19 fF in ND cells and 78±19 fF in T2D cells. This corresponds to a reduction of exocytosis by 25±10 % in T2D compared with ND (p=0.1, fig 1L), which is similar to the reduction observed by imaging granule release (Fig 1D). Cell size, as assessed by cell capacitance was not different in ND and T2D α-cells (Fig 1M).

### Intrinsic regulation of glucagon secretion

To study the glucose dependence of α-cell exocytosis, we imaged granule behavior in α-cells at steady-state conditions in different glucose concentrations (1, 3, 7, 10, or 20 mM, normal K^+^; Fig 2A-D), and quantified granule exocytosis (Fig 2B), docked granules (Fig 2C) and the rate of granule docking (Fig 2D). All three parameters displayed a bimodal relationship with the glucose concentration, with a nadir at 7 mM and a maximum at 20 mM glucose. Cells from T2D donors behaved essentially identically to cells from ND donors, with the same bimodal dose response to glucose and no significant differences at any of the tested concentrations. In both groups, the decrease in docked granules at 7 mM glucose, compared with 1 or 10 mM, was paralleled by reduced exocytosis (∼50%). Since exocytosis consumes docked granules (Fig 1C), the finding indicates that a reduced docking rate underlies the decrease in exocytosis at 7 mM glucose. Indeed, docking rates tended to be lower at 7 mM glucose compared with 1 or 10 mM (Fig 2D). To study the timecourse of the glucose dependent changes in exocytosis, we altered the glucose concentration stepwise, either between 10 and 7mM (Fig 2E) or between or between 1 and 7mM (Fig 2F). However, we were unable to detect changes in the exocytosis rate during the relatively short observation time, suggesting that isolated α-cells respond relatively slowly to changes in the ambient glucose concentration.

**Fig 2.**
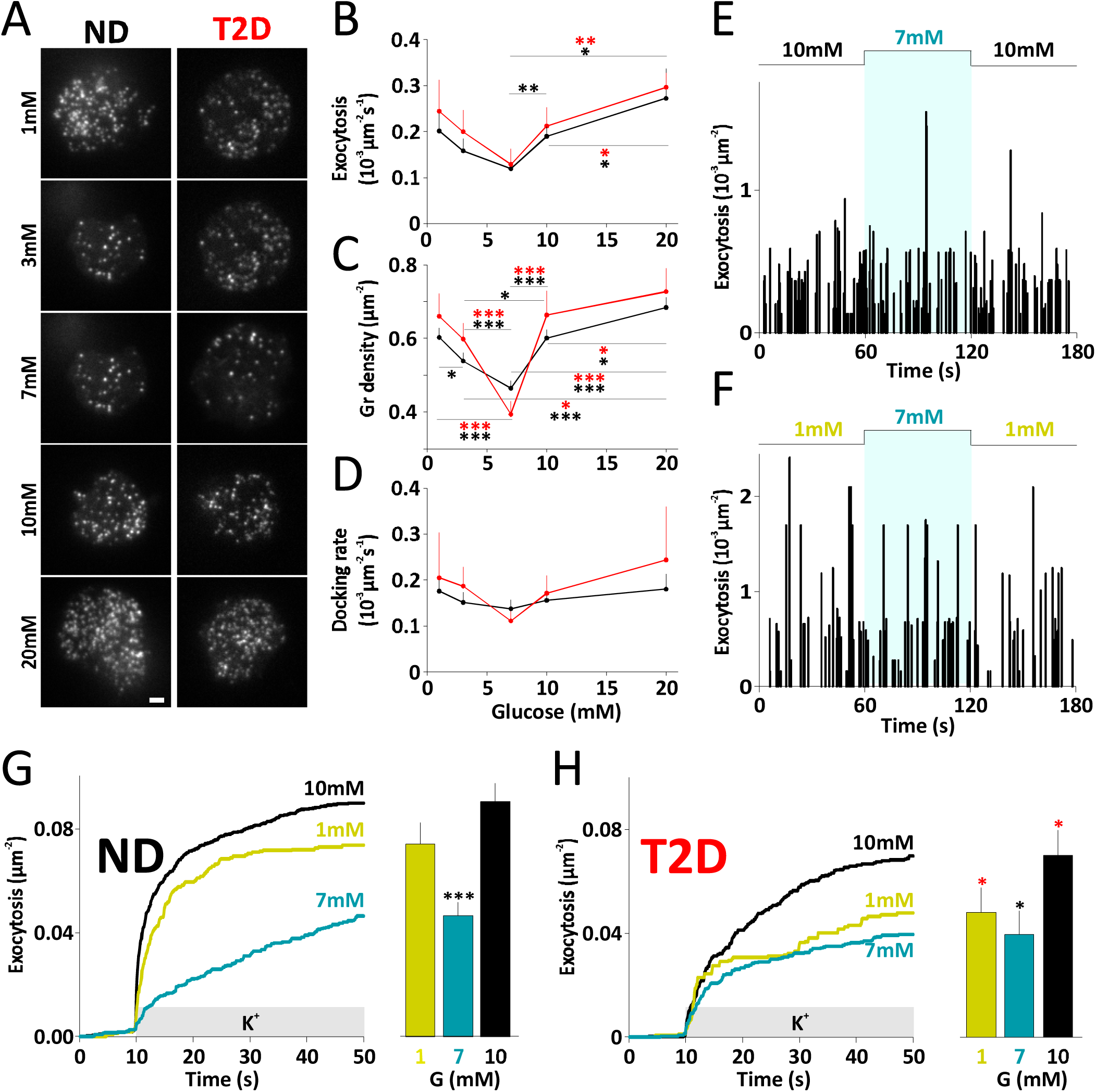
Different glucose levels exhibit different granule exocytosis profile in α-cells. **(A)** Examples TIRF images of non-diabetic (ND) cells (left) and type-2 diabetic (T2D) cells (right) expressing Pppg-NPY-EGFP, in presence of different glucose concentration: 1mM, 3mM, 7mM, 10mM or 20mM. Scale bar 2 µm. **(B)** Average exocytosis events in the TIRF field for ND cells (black) under different glucose conditions: 1mM (n=13 cells from 3 donors), 3mM (n=8 cells from 2 donors), 7mM (n=13 cells from 2 donors), 10mM (n=15 cells from 3 donors) or 20mM (n=10 cells from 2 donors), and for T2D cells (red) under different glucose conditions: 1mM (n=7 cells from 2 donors), 3mM (n=9 cells from 2 donors), 7mM (n=10 cells from 2 donors), 10mM (n=10 cells from 2 donors) or 20mM (n=7 cells from 2 donors). *;*; or *; indicates p<0.01 or p<0.05, respectively (t-test). Black or red asterisks indicate statistically significant differences between ND or T2D groups, respectively. **(C)** Average density of granules in the TIRF field for ND cells (black) under different glucose conditions: 1mM (n=32 cells from 5 donors), 3mM (n=32 cells from 5 donors), 7mM (n=32 cells from 5 donors), 10mM (n=32 cells from 5 donors) or 20mM (n=32 cells from 5 donors), and for T2D cells (red) under different glucose conditions: 1mM (n=11 cells from 2 donors), 3mM (n=11 cells from 2 donors), 7mM (n=17 cells from 2 donors), 10mM (n=16 cells from 3 donors) or 20mM (n=11 cells from 2 donors). *;*;*;, *;*; or *; indicates p<0.001, p<0.01 or p<0.05, respectively (t-test). Black or red asterisks indicate statistically significant differences between ND or T2D groups, respectively. **(D)** Average rate of docking in the TIRF field under different glucose conditions same like (B). **(E)** Plot of exocytosis frequency over time off cells exposed to 10 or 7mM glucose (blue bar). **(F)** Plot of exocytosis frequency over time off cells exposed to 1 or 7mM glucose (blue bar). **(G)** Time course of cumulative K^+^-stimulated exocytosis normalized to the cell area (K^+^ was elevated during 10-50 s, grey) for non-diabetic (ND) cells in the presence of different glucose concentration: 1mM (0.073±0.008 gr*;µm^-2^; n=16 cells from 3 donors, orange), 7mM (0.046±0.004 gr*;µm^-2^; n=28 cells from 4 donors, blue) and 10mM (0.089±0.007 gr*;µm-2; n=54 cells from 11 donors, black). Quantification of exocytosis (0-40 s) in experiments as in D (left). *;*;*; indicates p<0.001 (t-test). **(H)** As in G, but for diabetic (T2D) cells. *; indicates p<0.05 (t-test), Red asterisks indicate statistically significant differences in exocytosis between ND and T2D.

Next, we applied K^+^-stimulations to α-cells bathed in 1, 7, or 10 mM glucose (Fig 2G). This approach primarily elicits exocytosis of granules that are in a readily releasable state or “primed” for exocytosis. At all glucose concentrations, elevated K^+^ stimulated biphasic exocytosis that was rapid, compared with the relatively slow basal exocytosis. K^+^-stimulated exocytosis clearly followed a biphasic timecourse and was reduced by 37±10% or 47±7% in the presence of 7 mM, compared with both 1 and 10 mM glucose, respectively. This reduction was entirely due to a slower initial phase of exocytosis, which may correspond to the immediately releasable pool of granules in β-cells ^47^. In parallel, we observed a reduction in docked granules at 7 mM glucose, compared with both 1 and 10 mM (Fig S3 A), giving further support to the notion that the exocytotic capacity in α-cells is related to docked granules.

When α-cells from T2D donors were stimulated by depolarization with high K^+^, the rate of exocytosis was significantly lower than in ND α-cells (10 mM glucose, 0.069±0.009 gr*;µm^-2^; n=21 cells from 4 donors, p= 0.044, Fig 2H, black), and 1 mM condition (0.047±0.009 gr*;µm^-2^; n=12 cells from 2 donors, p= 0.025, Fig 2, yellow). 7 mM glucose had similar rate on T2D cells compared to ND cells (0.039±0.009 gr*;µm^-2^; n=24 cells from 3 donors, Fig 2H, blue). The density of docked granules was similar between ND and T2D cells in all three tested glucose concentrations (Fig S3). These data confirm that the secretory capacity of α-cells is diminished in type-2 diabetes, an effect at the level of granule priming that is most prominent in hypoglycemic conditions.

### Paracrine regulation of α-cell exocytosis

Glucagon secretion is regulated by a network of paracrine mechanisms, some of which act directly on α-cells. We therefore quantified K^+^-stimulated exocytosis in α-cells exposed to a panel of effectors somatostatin (SST), insulin (INS), forskolin (FSK), γ-aminobutyric acic (GABA), and adrenalin (ADR). As expected, the δ-cell hormone somatostatin inhibited K^+^-stimulated exocytosis by 65±5% (0.031±0.003 gr*;µm^-2^; n=41 cells from 7 donors, Fig 3A-B, blue), compared with control (10 mM glucose; 0.089±0.007 gr*;µm^-2^; n=54 cells from 11 donors, Fig 3A-B, black). Factors secreted by β-cells also affected α-cell exocytosis, with both insulin (0.04±0.003 gr*;µm^-2^; n=38 cells from 6 donors, Fig 3A-B, green) and GABA reducing it by about half (0.047±0.008 gr*;µm^-2^; n=16 cells from 3 donors, Fig 3A-B, brown). In contrast, adrenaline increased α-cell exocytosis by 46±27% (0.131±0.022 gr*;µm^-2^; n=23 cells from 3 donors, Fig 3A-B, pink). Elevated cAMP, after exposure to forskolin, had no effect on exocytosis (0.077±0.01 gr*;µm^-2^; n=30 cells from 5 donors, Fig 3A-B, purple), in contrast to previous reports ^50^. None of these compounds affected the density of docked granules (Fig 4C). This suggests that paracrine factors modulate α-cell exocytosis independently of granule docking, likely at the level of granule priming.

**Fig 3.**
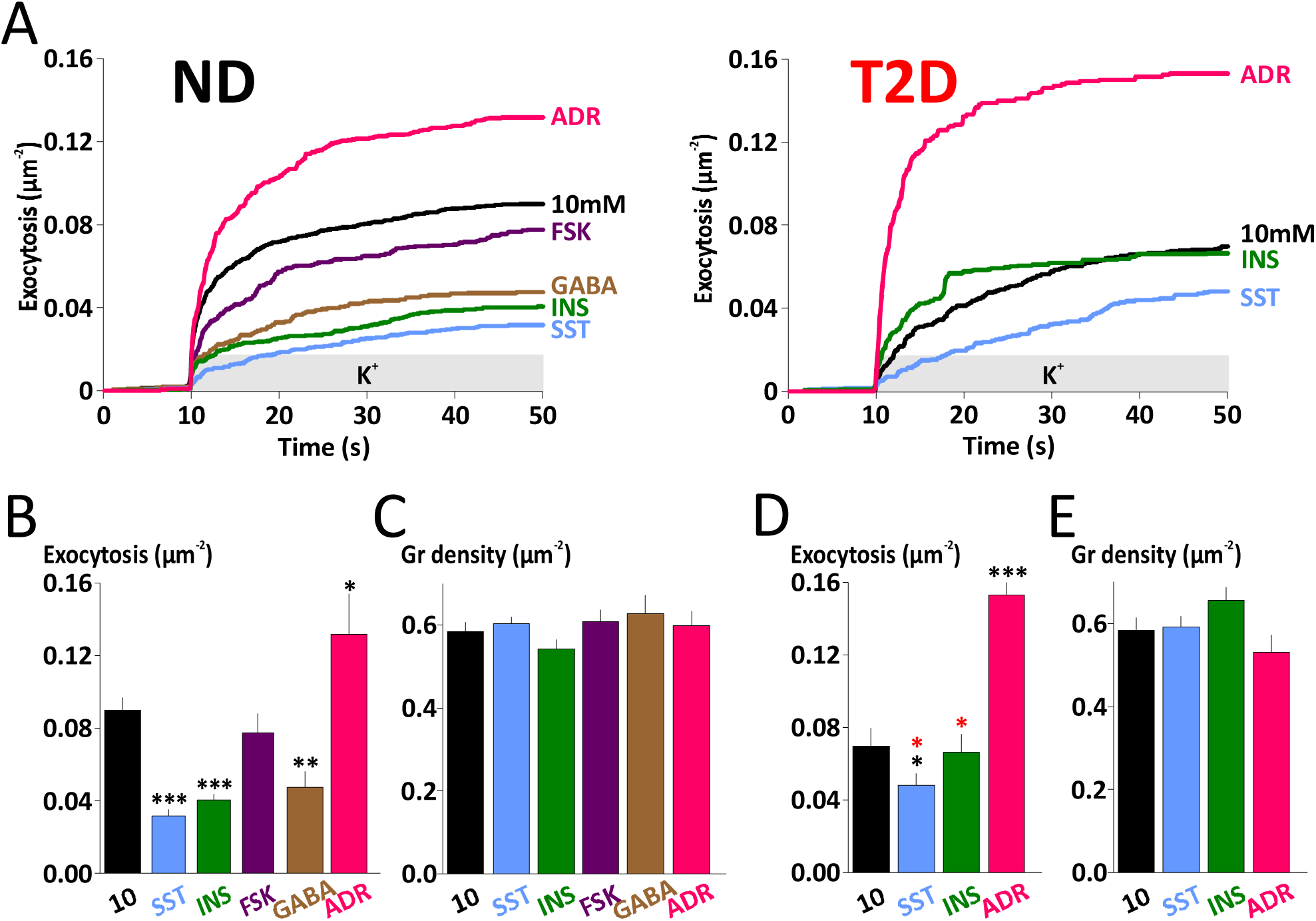
Paracrine regulation of glucagon secretion. **(A)** Average cumulative number of exocytotic events as function of time and normalized to the cell area (K+ was elevated during 10-50 s, grey) for non-diabetic (ND, left) or type-2 diabetic (T2D,right) cells exposed to adrenaline (ADR), forskolin (FSK), GABA, insulin (INS), somatostatin (SST), in the presence of 10mM glucose. **(B)** Total exocytosis after 40 s of stimulation in A (ND). *;*;*;, *;*; or *; indicates p<0.001, p<0.01 or p<0.05, respectively (t-test). **(C)** Average granule density normalized to footprint area of cells in the presence of different conditions as indicated (ND). **(D-E)** As in B-C, but for type-2 diabetic (T2D) cells. *;*;*;, *;*; or *; indicates p<0.001, p<0.01 or p<0.05, respectively (t-test).Red asterisks indicate statistically significant differences in exocytosis between ND and T2D.

**Fig 4.**
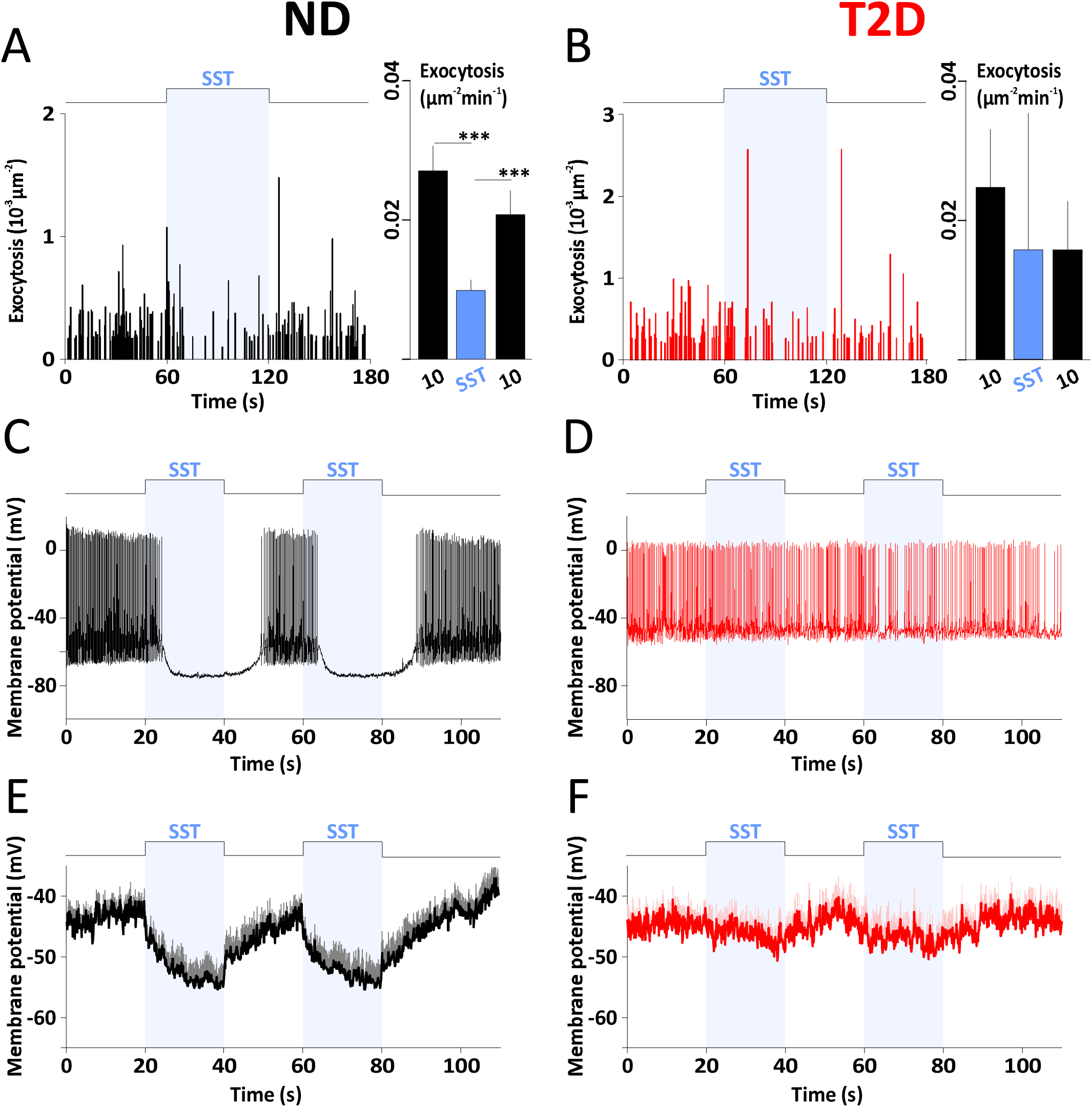
Somatostatin regulation of glucagon secretion. **(A)** Exocytosis events over time plotted for a representative cell from a ND donor exposed to 10 mM glucose and challenged with somatostatin (SST, 400 nM; left, blue bar). Average exocytosis for 36 cells from 6 donors during the periods indicated in the example (right). **(B)** Exocytosis events over time plotted for a representative cell from a T2D donor exposed to 10 mM glucose and challenged with somatostatin (SST, 400 nM; left, blue bar). Average exocytosis for 23 cells from 4 donors during the periods indicated in the example (right). **(C-D)** Examples of α-cells electrical activity from a ND or T2D donor in the presence of somatostatin (SST, 400 nM, blue bar). **(E-F)** Action potential frequency of α-cells from a ND (17 cells from 4 donors) or T2D (11 cells from 2 donors) donor in the presence of somatostatin (SST, 400 nM, blue bar).

In α-cells from T2D donors, exocytosis was also slowed by somatostatin (by 46±11%, 0.048±0.006 gr*;µm^-2^; n=20 cells from 5 donors, Fig 3A-D, blue), but this inhibitory effect was significantly less than in ND cells (p= 0.02). Notably, insulin-dependent inhibition of exocytosis was absent in T2D α-cells (0.066±0.009 gr*;µm-2; n=14 cells from 3 donors, p= 0.01 compared with ND cells, Fig 3A-D, green). Adrenaline accelerated exocytosis in T2D α-cells by 70±30% (0.15±0.016 gr*;µm^-2^; n=14 cells from 2 donors, Fig 3A-D, pink). No differences in the density of docked granules were observed in presence of any of the effectors, or compared with ND cells (Fig 3E, right).

Next, we determined the time course of paracrine inhibition of exocytosis by rapidly applying somatostatin to α-cells (Fig 4A, left). Maximal inhibition of exocytosis was reached within a few seconds (mono-exponential decay constant τ=9.4±5.3s n=36 cells, 6 donors), but frequently wore off towards the end of the 1 min challenge, and could be rapidly reversed by removing somatostatin (Fig 4A, right). Similarly, rapid effects on exocytosis were observed with with insulin (Fig S4A) or adrenaline (Fig S4B). Thus, paracrine regulation of glucagon exocytosis occurs on a timescale that is consistent with the observed glucagon pulsatility ^26^. Due to the small effect size, the time course of somatostatin inhibition of exocytosis could not be determined for T2D α-cells (Fig 4B).

Finally, we tested the effect of somatostatin on α-cell electrical activity by patch-clamp electrophysiology in the perforated patch configuration. In 10 mM glucose, electrical activity consisted of a train of action potentials as described previously ^39,51^. In ND α-cells, pulses of somatostatin (400 nM) applied to these cells caused rapid cessation of electrical activity (Fig 4C-D). In the example in Fig 4C, action potentials reappeared about 20 s after removal of the somatostatin. However, in many cells electrical activity returned already towards the end of the somatostatin pulse, which is apparent in the time course of the average action potential frequency (Fig 4E). Similar recovery during the somatostatin pulse was also seen for exocytosis (Fig 4A), and may reflect somatostatin receptor inactivation. In α-cells from T2D donors, the somatostatin pulse had little effect on the time course of exocytosis (Fig 4B) or electrical activity, which is consistent with the earlier conclusion that these cells are somewhat resistant to somatostatin.

## Discussion

Glucose controls glucagon secretion by intrinsic and paracrine mechanisms, but their relative significance is still debated ^52^ and secretory defects in type-2 diabetes are not well understood. The current work is first in using high resolution microscopy to study glucagon secretion in single, undisturbed α-cells of healthy and type-2 diabetic donors, thus isolating intrinsic from paracrine mechanisms while having full control over paracrine signaling. We show that in the absence of paracrine influence, isolated α-cells respond appropriately to hypoglycemia with an increase in glucagon granule exocytosis. This is consistent with the glucose dependence of glucagon secretion from intact islets (but not FACS sorted rat α-cells) ^52,53^, and indicates that glucagon secretion is mostly under intrinsic control in the lower glucose-concentration range. With only 2-fold difference in the exocytosis rate between minimal secretion at 7 mM and maximal secretion at 1 mM glucose, the dynamic range is small compared with β-cells. Notably, this glucose-dependent inhibition was absent in the hyperglycemic range, leading to a U-shaped response with nearly maximal exocytosis already at 10 mM glucose. This is in contrast to intact islets, in which glucagon secretion is depressed between 3 and 20 mM glucose, although even higher (non-physiological) concentrations of glucose can be stimulatory ^54^. Evidently, appropriate glucagon secretion in the hyperglycemic range is lost in isolated α-cells, and paracrine inhibition is likely responsible for the physiological response in this range. We did not observe any bursts of exocytosis, as might be expected given the pulsatile glucagon secretion from intact islets. This is in line with the absence of membrane potential oscillations in single cells ^55^ (that we confirm here), and indicates that the islet context is required not just for intra-islet synchronization, but for oscillatory α-cell behavior as such.

Elevated glucagon is a hallmark of type-2 diabetes ^56^. Despite this, both glucose-dependent and depolarization (K^+^)-induced exocytosis was reduced in α-cells from donors that had been diagnosed with T2D, and both exocytosis and docked granules were moderately anti-correlated with donor HbA1c values. This indicates that the exocytosis machinery in α-cells from type-2 diabetics is slightly impaired, while electrophysiological parameters (that determine electrical activity, depolarization and Ca^2+^-influx) were normal. The reason for this is unknown, but may reflect reduced expression of certain exocytosis-related proteins, as is the case in β-cells ^46,48,49^. The reduced exocytotic capacity in T2D α-cells was most apparent in hypo- and hyperglycemic conditions, i.e. when glucose was not inhibitory (Fig 2G-H). It is unrelated to glucose- or disease-related changes in electrical activity, because the membrane potential was clamped with elevated K^+^ during these experiments. Since exocytosis in single α-cells is impaired rather than increased in T2D, the hyperglucagonemia in diabetic humans must be due to mechanisms that are lost in isolated cells, such as paracrine or neuronal regulation ^57^. In addition, gut derived glucagon may contribute to hyperglucagonemia following oral glucose intake ^58,59^.

Exposure to insulin, somatostatin, and GABA reduced α-cell exocytosis, while adrenalin stimulated it. This is consistent with the known effects of these signaling molecules on islets, as well as systemically ^52^. We show here for somatostatin that these effects are very rapid (seconds), which is consistent with the frequency of pulsatile glucagon release in vivo and from intact islets. In the absence of glucose-dependent control, paracrine inhibition by insulin and somatostatin is therefore the most likely mechanism for glucagon regulation in the hyperglycemic range. It can be speculated that the differential glucose dependence of insulin and somatostatin secretion is reflected in different target glucose ranges for their action on α-cells. Strikingly, the inhibitory effect of insulin and somatostatin on glucagon exocytosis was strongly reduced in cells from T2D donors. Together with the overall reduced exocytosis, this points to α-cell resistance to insulin and somatostatin as the main cause for inadequate glucagon secretion in type-2 diabetes, which in turn exacerbates hyperglycemia ^60^. Insulin resistance is a hallmark of T2DM, and has previously been proposed as mechanism for hyperglucagonemia. For example, insulin resistance is associated with fasting glucagon levels ^38^, and this inverse relationship is lost in type-2 diabetes ^28^. All tested paracrine modulators affected exocytosis machinery at the priming step, rather than by increasing granule docking. This is consistent with previous findings that somatostatin inhibits exocytosis in rat α-cells through G_i_-dependent depriming ^32^, and reports that antagonists of SSTR2 ^12^ or the associated G-protein cascade ^12^ increase glucagon secretion without altering the glucose-dependent inhibition of glucagon secretion.

Capacitance measurements indicate that glucagon granules exist in at least two states with different release probabilities, which are often referred to as the readily releasable pool of granules (RRP) and a larger reserve pool (RP) ^42,61,62^. We show here that glucagon granules were present at the plasma membrane for extended periods before undergoing exocytosis. We interpret this as the relatively slow conversion from RP to RRP that reflects the molecular assembly of the secretory machinery at the release site, in analogy with the situation in β-cells ^44,46^. Throughout the glucose range, the rate of exocytosis was nearly identical to that of granule docking (Fig 1B-D), suggesting that docking is rate limiting for secretion. This may indeed be the case during strong (non-physiological) stimulation, as illustrated by the finding that the glucose-dependence of depolarization-induced exocytosis followed that of granule docking. However, in physiological conditions elevated K^+^ accelerated exocytosis about 50-fold (during the first second), which indicates a large excess in exocytotic capacity that is not triggered by normal α-cell electrical activity. A possible explanation could be that only a limited number of granules is positionally primed, i.e. located near voltage-gated Ca^2+^-granules ^63^. Further theoretical work is required to understand the combination of factors affecting granule exocytosis, and the granule conversion rates provided here may be useful in this regard.

## Acknowledgment

We thank Jan Saras for expert technical assistance, and Hongyan Shuai and Yunjian Xu for supplying Pppg-GFP plasmid and adenovirus. Human islets were provided through the JDRF award 31-2008-416 (ECIT Islet for Basic Research Program) and the Alberta Diabetes Institute Islet-Core, University of Alberta (with assistance of the Human Organ Procurement and Exchange [HOPE] program, Trillium Gift of Life Network [TGLN], and other Canadian organ procurement organizations). The work was supported by the Swedish Science Council, Diabetes Wellness Network Sweden, the Swedish Diabetes Society, the European Foundation for the Study of Diabetes, and the NovoNordisk Foundation, Excellence of Diabetes Research in Sweden (EXODIAB), and the Family Ernfors- and OE&E Johanssons-Foundations.

## Author contributions

M.O.H. and S.B. designed experiments and analyzed the data. M.O.H. performed experiments. P.E.L. performed electro-physiology experiments. M.O.H. and N.R.G. isolated and picked human islets. M.O.H. designed and generated the Pppg-NPY-GFP virus construct. A.T supplied Pppg-GFP plasmid and adenovirus. S.B. conceived the study. S.B. and M.O.H. wrote, and all authors critically reviewed the manuscript.

## Declaration of Interests

The authors declare no competing interests.

## Online Methods

### Cells

Human pancreatic islets were obtained from the Nordic Network for Clinical Islet Transplantation Uppsala ^64^ or the ADI Isletcore at the University of Alberta ^65^ under full ethical clearance (Uppsala Regional Ethics Board 2006/348 and Alberta Human Research Ethics Board, Pro00001754) and with the donor families’ written informed consent. Isolated islets were cultured free-floating in sterile dishes in CMRL 1066 culture medium containing 5.5mM glucose, 10% fetal calf serum (FCS), 2mM L-glutamine, streptomycin (100U/ml), and penicillin (100U/ml) at 37°C in an atmosphere of 5% CO_2_ up to two weeks. Before imaging, islets were dispersed into single cells in 2 mL cell dissociation buffer (Thermo Fisher Scientific) supplemented with trypsin (0.005%, Life Technologies) and then gentle pipetting for 30 seconds. The trypsin was then inhibited by adding four mL serum-containing medium and centrifuged for 5 minutes at 160 g. Cells plated onto 22-mm polylysine-coated coverslips and allowed to settle overnight and infected then with different adenoviruses.

### Labeling of human pancreatic α-cells and glucagon granules

To identify α-cells, we transduced dispersed human islets with adenovirus coding for enhanced green fluorescent protein (EGFP) under the control of the pre-proglucagon promoter (Shuai et al., 2016). The system takes advantage of Tet-On conditional expression in the presence of 4 μM doxycycline, to drive expression of sufficient amounts of EGFP in spite of the relatively weak pre-proglucagon promoter. The cells were simultaneously transduced with adNPY-mCherry, a well-established secretory granule marker. Alternatively, α-cells granules were labeled with adenovirus coding for EGFP-tagged neuropeptide Y under control of the pre-proglucagon promoter (Pppg-NPY-EGFP), thus using only one construct to identify cell type and label secretory granules. For both approaches, immunostaining with an anti-glucagon antibody confirmed that over 90% of the fluorescently labeled cells were α-cells (Fig 1A). Confocal imaging of the immunostained cells indicated that approximately one-third of the glucagon positive cells were also labeled with Pppg-NPY-EGFP (Fig S1A). NPY-EGFP labeled granules had excellent overlap with punctate glucagon staining (Fig 1A), with 94±1% of glucagon positive granules being labeled with NPY-EGFP (37 cells, 5 donors). Thus, genetic labelling allows simple and robust identification of α-cells within pancreatic cell populations, which can be used for live-cell imaging or electrophysiology. We verified that exocytosis in the identified cells was strongly stimulated by adrenaline (Fig S1B-D), which increases intracellular Ca^2+^ in α-but not β-cells. The rate was slightly slower than with elevated K^+^ (0.058±0.015 gr*;µm^-2^; n=19 cells from 3 donors, Fig S1D, blue).

### Dual pulse stimulation protocol

We designed a protocol with two successive K^+^ depolarizations (40 s each) and separated by a 2 min rest period, and quantified the recovery of exocytosis and docked granules. The second depolarization tested to what extent the remaining or newly docked granules had acquired release competence during the resting interval.

### Solutions

Cells were imaged in a standard solution containing (in mM) 138 NaCl, 5.6 KCl, 1.2 MgCl_2_, 2.6 CaCl_2_, 10 D-glucose, 5 HEPES (pH 7.4 with NaOH). In Fig 2, the D-glucose concentration was varied (1 mM, 3 mM, 7 mM, 10 mM and 20 mM). Exocytosis was evoked with high K^+^ solution (75 mM KCl equimolarly replacing NaCl in the standard solution). High K^+^ solution was applied by computer-timed local pressure ejection from a pulled glass pipette (similar to those used for patch clamp). In Fig 3,4, cells were imaged in standard solution (10 D-glucose) in the presence of forskolin (Fsk; 2 µM), somatostatin (SST; 400 nM), insulin (INS; 100 nM), GABA (400 nM) or adrenaline (ADR; 5 µM).

### Microscopy

Cells were imaged using a custom-built lens-type total internal reflection (TIRF) microscope based on an AxioObserver Z1 with a 100x/1.45 objective (Carl Zeiss). Excitation was from two DPSS lasers at 491 and 561 nm (Cobolt) passed through a cleanup filter (zet405/488/561/640x, Chroma) and controlled with an acousto-optical tunable filter (AA-Opto). Excitation and emission light were separated using a beamsplitter (ZT405/488/561/640rpc, Chroma). The emission light was chromatically separated onto separate areas of an EMCCD camera (Roper QuantEM 512SC) using an image splitter (Optical Insights) with a cutoff at 565 nm (565dcxr, Chroma) and emission filters (ET525/50m and 600/50m, Chroma). Scaling was 160 nm per pixel.

Confocal microscopy was done with a Zeiss LSM780 using a 63/1.40 objective (Zeiss) with sequential scanning of the red (excitation 561 nm, emission 578–696 nm) and green channel (excitation 488 nm, emission 493–574 nm). Pinhole size was 0.61 mm, corresponding to 1 Airy unit. Images were acquired in 16-bit at gain settings 750 for both channels and 0.11 mm per pixel.

### Image analysis

Exocytosis events were identified based on the characteristic rapid loss of the granule marker fluorescence (1-2 frames). Granule docking events were rare and defined as granules that approached the TIRF field and becoming laterally confined once they reached their maximum brightness ^66^.

Docked granules were counted using the ‘find maxima’ function in ImageJ (http://rsbweb.nih.gov/ij). Values were normalized to each cells’ contact area with the coverslip (footprint). The *ΔF* parameter estimates the fluorescence that is specifically localized to a granule, but subtracting a local background value (average of a 5 pixel wide annulus) from the average fluorescence value in a 3 pixel wide circle, both centered at the granule position.

### Electrophysiology

Standard whole-cell voltage clamp and capacitance recordings were performed using an EPC-9 patch amplifier (HEKA Electronics, Lambrecht/Pfalz, Germany) and PatchMaster software. Voltage-dependent currents were investigated using an IV-protocol, in which the membrane was depolarized from −70 mV to +80 mV (10 mV steps) during 50 ms each. Currents were compensated for capacitive transients and linear leak using a *P/*4 protocol. Exocytosis was detected as changes in cell capacitance using the lock-in module of Patchmaster (30 mV peak-to-peak with a frequency of 1 kHz).

Patch electrodes were made from borosilicate glass capillaries coated with Sylgard close to the tips and fire-polished. The pipette resistance ranged between 2 and 4 MΩ when filled with the intracellular solution containing (in mM) 125 Cs-glutamate, 10 CsCl, 10 NaCl, 1 MgCl_2_, 0.05 EGTA, 3 Mg-ATP, 0.1 cAMP, and 5 HEPES, pH 7.2 adjusted using CsOH.

During the experiments, the cells were continuously superfused with an extracellular solution containing (in mM) 138 NaCl, 5.6 KCl, 1.2 MgCl_2_, 2.6 CaCl_2_, 10 D-glucose, and 5 Hepes, pH 7.4 adjusted with NaOH at a rate of 0.4 ml/min. All electrophysiological measurements were performed at 32 C. In the analysis of the measured voltage-dependent current consists of both Na^+^ and Ca^2+^ current components, were the rapid peak current (0-3 ms) represent the Na^+^ current and the sustained current during the latter part of the depolarization reflects the Ca^2+^ current. In Fig4, for membrane potential, solution was containing (in mM) 76 K_2_SO_4_, 10 KCl, 1 MgCl_2_and 5 HEPES, pH 7.3 adjusted with KOH.

### Statistics

Data are presented as mean ± SEM unless otherwise stated. Statistical significance was tested using Students t-test and is indicated by asterisks (*;p < 0.05, *;*;p < 0.01, *;*;*;p < 0.001). Correlation was quantified as Pearson coefficient r using Excel.

## Supplementary legends

**Fig S1.**
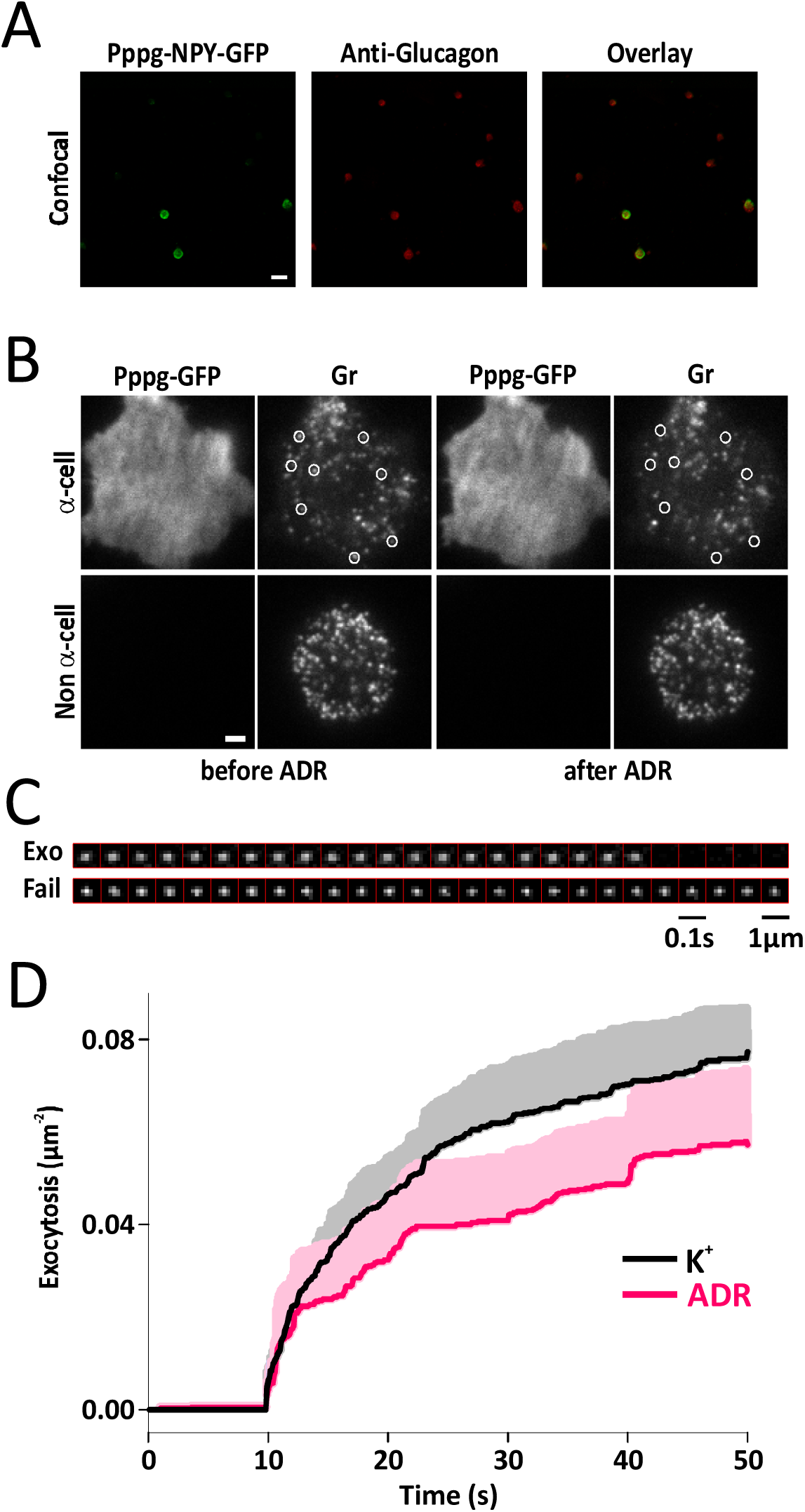
Identification of human pancreatic α-cells. (**A**) Confocal images of Pppg-NPY-EGFP expressing and glucagon positive cells in a dispersed human islet α-cells. Scale bar 20 µm. (**B**) TIRF images of α-cells (top) expressing Pppg-EGFP together with NPY-Cherry and non α-cells (bottom) expressing NPY-Cherry before and after stimulation with 5µM adrenaline for 40s. Scale bar 2 µm. (**C**) Image sequences showing example of exocytotic (exo, top) and failure (fail, bottom) event during stimulation of cells as in (**B**). (**D**) Average cumulative number of exocytotic events in adrenaline stimulation experiments (pink) or K^+^ stimulation experiments (black) plotted over time and normalized to the cell area.

**Fig S2.**
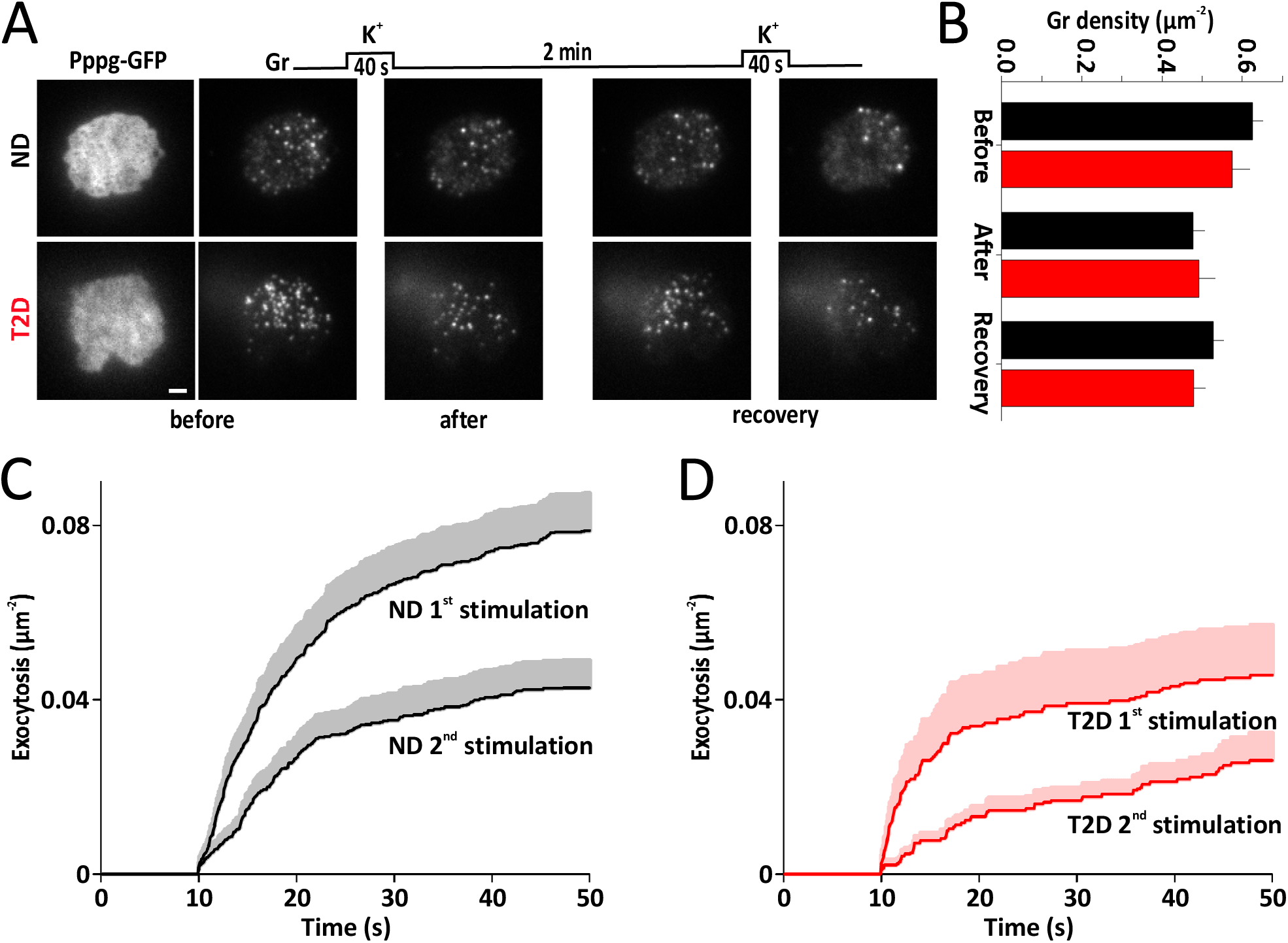
Depletion and recovery of docked granules during K^+^ induced exocytosis. (**A**) Examples of ND cells (top) and T2D cells (bottom) at different times during a dual pulse stimulation protocol. Scale bar 2 µm. (**B**) Average granule density of cells from ND (black) and T2D donors (red) at different times during a dual pulse stimulation protocol. The initial density of docked granules in T2D α-cells was 0.57 gr*;µm-2 compared with 0.62 gr*;µm-2 in cells of ND donors (p=NS; 7 ND vs 3 T2D donors with 5-15 cells each). The first depolarization evoked exocytosis of about 15% of the docked granules in T2D α-cells (n=17) and 24 % of the docked granules in ND cells (n=55; p<n.s). After the 2 min resting period, recovery of docked granules was negligible in T2D α-cells, compared with 8% in ND α-cells. (**C**) Average cumulative number of exocytotic events in ND experiments as in (A) plotted over time and normalized to the cell area for first and second stimulation. Stimulation after the 2 min rest period indicated partial recovery of the exocytotic capacity in ND α-cells (58% of initial). (**D**) Average cumulative number of exocytotic events in T2D experiments as in (A) plotted over time and normalized to the cell area for first and second stimulation. Stimulation after the 2 min rest period indicated partial recovery of the exocytotic capacity in both T2D α-cells (54% of initial).

**Fig S3.**
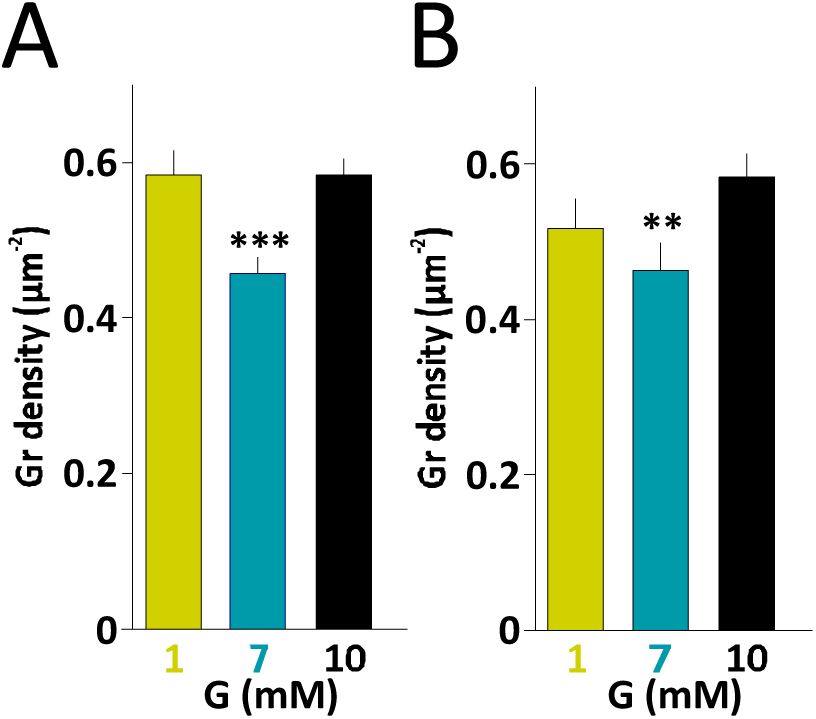
Different glucose levels exhibit different granule density. **(A)** Average granule density normalized to footprint area of cells in the presence of different conditions as indicated **(B)** As in B, but for diabetic (T2D) cells.

**Fig S4.**
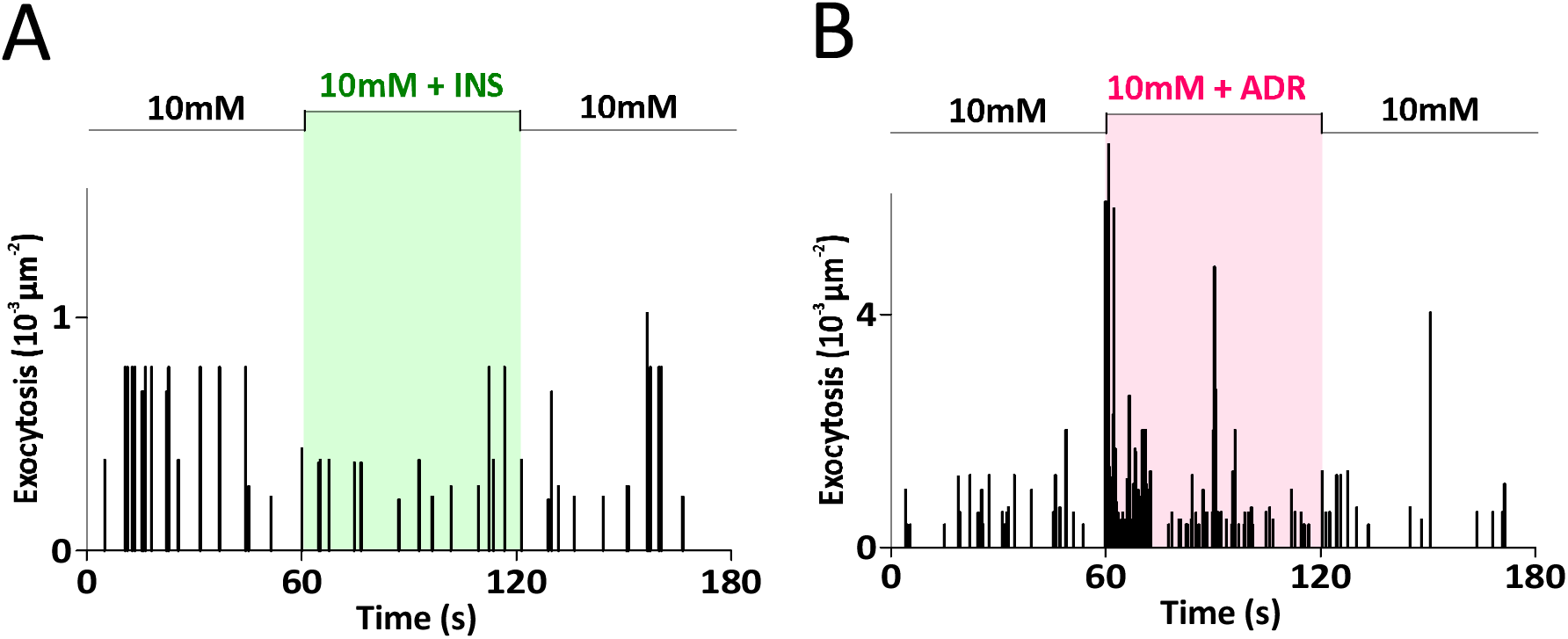
Regulation of glucagon exocytosis by insulin and adrenaline. (**A**) Plot of exocytosis events over time of cells exposed to 10mM glucose in the presence or absence of insulin. (**B**) Plot of exocytosis events over time of cells exposed to 10mM glucose in presence or absence of adrenaline.

**Table S1.**
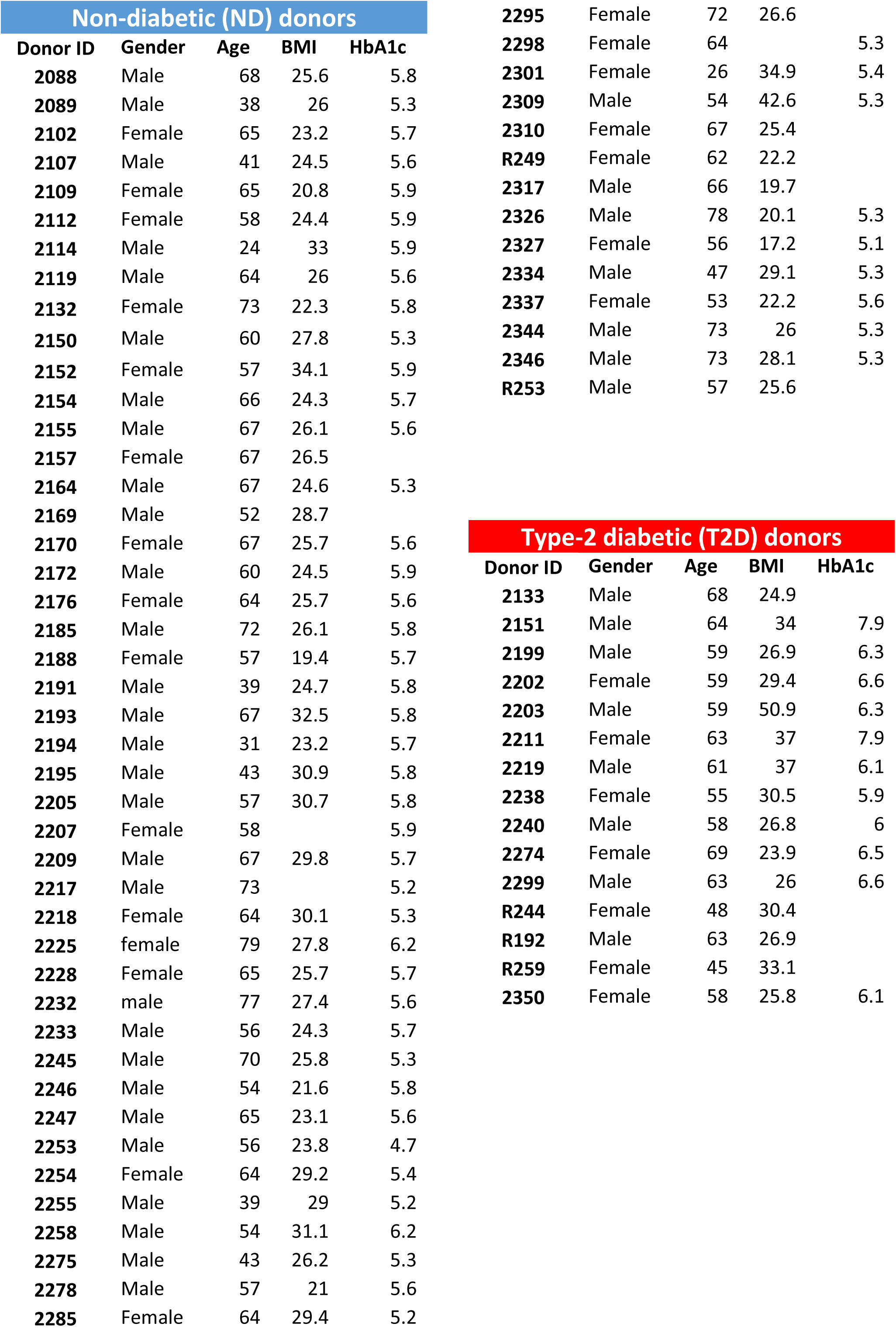
Human donor information.

**Fig S4**
| Non-diabetic (ND) donors |  |  |  |  |
| --- | --- | --- | --- | --- |
| Donor ID | Gender | Age | BMI | HbA1c |
| 2088 | Male | 68 | 25.6 | 5.8 |
| 2089 | Male | 38 | 26 | 5.3 |
| 2102 | Female | 65 | 23.2 | 5.7 |
| 2107 | Male | 41 | 24.5 | 5.6 |
| 2109 | Female | 65 | 20.8 | 5.9 |
| 2112 | Female | 58 | 24.4 | 5.9 |
| 2114 | Male | 24 | 33 | 5.9 |
| 2119 | Male | 64 | 26 | 5.6 |
| 2132 | Female | 73 | 22.3 | 5.8 |
| 2150 | Male | 60 | 27.8 | 5.3 |
| 2152 | Female | 57 | 34.1 | 5.9 |
| 2154 | Male | 66 | 24.3 | 5.7 |
| 2155 | Male | 67 | 26.1 | 5.6 |
| 2157 | Female | 67 | 26.5 |  |
| 2164 | Male | 67 | 24.6 | 5.3 |
| 2169 | Male | 52 | 28.7 |  |
| 2170 | Female | 67 | 25.7 | 5.6 |
| 2172 | Male | 60 | 24.5 | 5.9 |
| 2176 | Female | 64 | 25.7 | 5.6 |
| 2185 | Male | 72 | 26.1 | 5.8 |
| 2188 | Female | 57 | 19.4 | 5.7 |
| 2191 | Male | 39 | 24.7 | 5.8 |
| 2193 | Male | 67 | 32.5 | 5.8 |
| 2194 | Male | 31 | 23.2 | 5.7 |
| 2195 | Male | 43 | 30.9 | 5.8 |
| 2205 | Male | 57 | 30.7 | 5.8 |
| 2207 | Female | 58 |  | 5.9 |
| 2209 | Male | 67 | 29.8 | 5.7 |
| 2217 | Male | 73 |  | 5.2 |
| 2218 | Female | 64 | 30.1 | 5.3 |
| 2225 | female | 79 | 27.8 | 6.2 |
| 2228 | Female | 65 | 25.7 | 5.7 |
| 2232 | male | 77 | 27.4 | 5.6 |
| 2233 | Male | 56 | 24.3 | 5.7 |
| 2245 | Male | 70 | 25.8 | 5.3 |
| 2246 | Male | 54 | 21.6 | 5.8 |
| 2247 | Male | 65 | 23.1 | 5.6 |
| 2253 | Male | 56 | 23.8 | 4.7 |
| 2254 | Female | 64 | 29.2 | 5.4 |
| 2255 | Male | 39 | 29 | 5.2 |
| 2258 | Male | 54 | 31.1 | 6.2 |
| 2275 | Male | 43 | 26.2 | 5.3 |
| 2278 | Male | 57 | 21 | 5.6 |
| 2285 | Female | 64 | 29.4 | 5.2 |
| 2295 | Female | 72 | 26.6 |  |
| 2298 | Female | 64 |  | 5.3 |
| 2301 | Female | 26 | 34.9 | 5.4 |
| 2309 | Male | 54 | 42.6 | 5.3 |
| 2310 | Female | 67 | 25.4 |  |
| R249 | Female | 62 | 22.2 |  |
| 2317 | Male | 66 | 19.7 |  |
| 2326 | Male | 78 | 20.1 | 5.3 |
| 2327 | Female | 56 | 17.2 | 5.1 |
| 2334 | Male | 47 | 29.1 | 5.3 |
| 2337 | Female | 53 | 22.2 | 5.6 |
| 2344 | Male | 73 | 26 | 5.3 |
| 2346 | Male | 73 | 28.1 | 5.3 |
| R253 | Male | 57 | 25.6 |  |

**Table S1 – Human donor information**
| Type-2 diabetic (T2D) donors |  |  |  |  |
| --- | --- | --- | --- | --- |
| Donor ID | Gender | Age | BMI | HbA1c |
| 2133 | Male | 68 | 24.9 |  |
| 2151 | Male | 64 | 34 | 7.9 |
| 2199 | Male | 59 | 26.9 | 6.3 |
| 2202 | Female | 59 | 29.4 | 6.6 |
| 2203 | Male | 59 | 50.9 | 6.3 |
| 2211 | Female | 63 | 37 | 7.9 |
| 2219 | Male | 61 | 37 | 6.1 |
| 2238 | Female | 55 | 30.5 | 5.9 |
| 2240 | Male | 58 | 26.8 | 6 |
| 2274 | Female | 69 | 23.9 | 6.5 |
| 2299 | Male | 63 | 26 | 6.6 |
| R244 | Female | 48 | 30.4 |  |
| R192 | Male | 63 | 26.9 |  |
| R259 | Female | 45 | 33.1 |  |
| 2350 | Female | 58 | 25.8 | 6.1 |

